# eQTLs are key players in the integration of genomic and transcriptomic data for phenotype prediction

**DOI:** 10.1101/2021.09.07.459279

**Authors:** Abdou Rahmane Wade, Harold Duruflé, Leopoldo Sanchez, Vincent Segura

## Abstract

Multi-omics represent a promising link between phenotypes and genome variation. Few studies yet address their integration to understand genetic architecture and improve predictability. Our study used 241 poplar genotypes, phenotyped in two common gardens, with their xylem and cambium RNA sequenced at one site, yielding large phenotypic, genomic and transcriptomic datasets. For each trait, prediction models were built with genotypic or transcriptomic data and compared to concatenation integrating both omics. The advantage of integration varied across traits and, to understand such differences, we made an eQTL analysis to characterize the interplay between the genome and the transcriptome and classify the predicting features into CIS or TRANS relationships. A strong and significant negative correlation was found between the change in predictability and the change in predictor importance for eQTLs (both TRANS and CIS effects) and CIS regulated transcripts, and mostly for traits showing beneficial integration and evaluated in the site of transcriptomic sampling. Consequently, beneficial integration happens when redundancy of predictors is decreased, leaving the stage to other less prominent but complementary predictors. An additional GO enrichment analysis appeared to corroborate such statistical output. To our knowledge, this is a novel finding delineating a promising way to explore data integration.

**One-sentence summary:** Successful multi-omics integration when predicting phenotypes makes redundant the predictors that are linked to ubiquitous connections between the omics, according to biological and statistical approaches

## Introduction

Genomic prediction, the prediction of phenotypes with genome-wide polymorphisms, has become a key tool to plant and animal breeders. This approach relies on statistical modelling to infer the effect of genomic variants, with many different modeling alternatives proposed in the literature (de los Campos et al., 2013; Varona et al., 2018). These models are mostly devised to predict the additive and transmissible contribution to individual genetic values, although dominance and epistatic interactions can also be accounted for Varona et al. (2018). Despite their success in identifying relevant effects and predicting phenotypes accurately, even in their most complex formulations, these models do not capture *per se* the genetic architecture of complex traits (Gianola, 2021). Beyond the statistics, it is the use of biological and functional information from the different organizational layers lying between the raw sequence and the organismal phenotype that will likely provide the required insights to reveal genetic architectures. Layers such as DNA methylation (Epigenome), transcripts (Transcriptome), proteins (Proteomics) or metabolites (Metabolites), are nowadays becoming increasingly accessible for many species, opening prospects towards a better understanding of the genetic architecture of complex traits.

In order to simultaneously account for these different layers of data in phenotype prediction, several integration approaches have been proposed (Ritchie et al., 2015). Among those, the most frequently used approach is the transformation or kernel-based integration, which consists in transforming each omics data into an intermediate form, usually taking the shape of a relationship matrix between the individuals (Guo et al., 2016; Schrag et al., 2018; Li et al., 2019; Morgante et al., 2020). Effects owing to different omics can then be integrated into a single analytical model, each effect being associated to a given kernel. Eventually, different kernels can be further combined by Hadamard product to add extra interaction terms between effects (Guo et al., 2016; Morgante et al., 2020). Integration can also be carried out across models, in what is known as model-based integration (Ritchie et al., 2015). Such integration can happen for a given omic type over different datasets or populations, each one summarized by its own model, with a final global model feeding on the top features contributed by each of the initial models. Another variant of the same model-based integration proceeds through a multistage approach, combining sequentially different omics for a given population (Ye et al., 2020). One of the simplest integration approaches, however, remains data concatenation (Azodi et al., 2020), by which multiple omics are placed side by side into a single large input matrix. Unlike kernels, whose results are produced at the individual level, the concatenation approach allows for the effects of multiple features at each omic to be estimated, whether they are SNPs, transcripts or any other omic. Another advantage, derived from that atomization of effects, is the fact that interactions between omics can be more easily captured, without the risk of being lost by intermediate transformations.

Most of the studies dealing with omics integration for phenotypic prediction have focused on gauging predictive abilities. To that level, the reported benefits of integration are context dependent across studies and, in general, amounting to small differences when compared to single omics counterparts. A series of published comparisons in maize illustrates this point. Using kernels to integrate genomic and transcriptomic data, Guo et al. (2016) found improved accuracies over single omic counterparts for most of the 11 economically important traits under study. Schrag et al. (2018), on the contrary, found no benefit in integration following a similar approach and on a similar set of production-related traits. For Azodi et al. (2020), however, using concatenation of genomic and transcriptomic data for three maize traits yielded benefits only for one of the traits. Studies on other biological models also showed similar context dependent results. Based on the Drosophila melanogaster Reference Panel and different transcriptomic datasets, Li et al. (2019) and Morgante et al. (2020) found subsequently no benefit of integration following a multiple kernel approach in terms of predictive abilities, and over different sets of fitness-related traits. When the integration included a gene ontology (GO) category as an additional layer of information, accuracies were surprisingly improved (Morgante et al., 2020). Using the same Drosophila panel, however, Ye et al. (2020) found some benefits by following a model-based integration approach, with a first modeling stage aiming at detecting SNP associated with transcripts (eQTLs), and a subsequent prediction model focused on resulting eQTLs. The number of studies, however, is not yet high enough to draw general conclusions. Benefits might depend jointly on methods of integration and targeted traits, reflecting the complexity of underlying architectures and conditions of studied populations.

Beyond the reported differences in prediction performance, there is still a scarcer number of studies available that were able to pinpoint some of the possible causes underlying the changes brought by integration to prediction. Already, at statistical level, omics like sequence polymorphisms and transcriptomics are likely non-orthogonal to some extent. If such redundancy is not conveniently handled at the model level, one can expect inaccurate estimation of effects and impaired prediction accuracy as a result (Farrar and Glauber, 1967; Ritchie et al., 2015). Redundancy between genomic and transcriptomic data was addressed in a few studies, typically by gauging the amount of extra variance captured by the different integration models compared to single omic counterparts. For instance, the successful integration described by Guo et al. (2016) was systematically accompanied by extra levels of captured variance, suggesting that each extra layer added to the model contributed to some extent with non-redundant information, and thus improving the prediction. The opposite behavior is described in Morgante et al. (2020), with integrative models showing similar levels of captured variance to those of single omics, indicating high levels of redundancy. It is interesting to note here that for this latter study, redundancy was not found between GO terms, the only layer bringing benefits to integration in the study. The most explanatory GO terms with genomic data were different from those detected for transcriptomic data. The second, more biological, approach is to look to what extent the most important features in both omics show at the same time mutual associations, in other words, if relevant SNPs are associated or not to relevant transcripts for a given phenotype. Azodi et al. (2020) showed, in maize, that the transcriptome brings information on the phenotype that is different from the one brought by genomic polymorphisms, by highlighting that the information carried by the most important transcripts to predict flowering time is not redundant with that carried out by the most important SNPs. In mice, two independent studies used a Bayesian approach to model the phenotype with both genomic and transcriptomic data (Ehsani et al., 2012; Takagi et al., 2014). These studies showed that specific SNPs (eQTLs) associated with gene expression profiles can contribute to the observed redundancy between the two data sources, which is reflected by the fact that their importance for phenotype prediction was substantially affected under the integrative approach.

Further research is needed to enrich the number of studies in data integration. It is clear that the mechanisms by which integration is successful when predicting phenotypes are still not known precisely and over a wide range of conditions and species, with the hypothesis of redundancy being one of the possible explanations. To some extent, redundancy reflects interactivity in the highly integrative space going from the raw genomic sequence to the organismal phenotype. Both redundancy and interactivity are key features to understand genetic architecture beyond the simple list of effects that is typically provided by genomic approaches. Most of the available studies on data integration involve model species like drosophila, maize, and notably humans. In the present study, we proposed new insights on data integration for a species not frequently found as subject of these approaches, black poplar, and using one of the simplest integration alternatives (concatenation) combined to one of the most popular prediction approaches (ridge regression). Here, we aimed at evaluating the factors affecting prediction accuracy when integrating genomic and transcriptomic data for phenotype prediction. Using a fairly large number of diverse phenotypes collected in two common gardens for a collection of black poplars, we specifically analyzed the change in importance of each of the potentially redundant sources (eQTLs and their target genes) between a multi-omics model and the single omics counterparts, together with the evolution of prediction accuracy. Under a more functional point of view, we further studied the redundancy using a Gene ontology (GO) enrichment analysis.

## Results

### Multi-omics model displays performance advantages over the single omic ones for specific functional types of traits

Twenty-one traits of different types were phenotyped (**Table 1**) from 241 poplars grown in two common garden experiments located at contrasting sites (Orleans, France, and Savigliano, Italy). RNA sequencing analysis was also performed on young differentiating xylem and cambium tissues of the entire set of genotypes sampled in the common garden located at Orleans, resulting in large genomic (428,836 SNPs) and transcriptomic (34,229 transcripts) datasets. For each phenotypic trait, three ridge regression models were built: the first one with genotypic data as predictors (denoted G), the second one with transcriptomic data as predictors (T), and the third one with integration by concatenation of both omics data (G+T). The prediction accuracies for the three models varied across trait types, with growth, pathogen tolerance and phenology traits having average performances above 0.5 on both testing sites, while biochemical and architectural traits having average performances below 0.5 (**Figure 1**).

**Table 1:** List of phenotypic traits. List of phenotypic traits used in the study with their abbreviations, classified by functional types, with site of measurement and year.

| Functional types | Trait | Abbreviation | Site | Year |
| --- | --- | --- | --- | --- |
| Growth | Height | Ht | ORL | 2011 |
|  | Circumference | Circ | ORL | 2011 |
|  |  |  | SAV | 2009 |
| Pathogen Tolerance | Tolerance to rust | Rust | ORL | 2009 |
| Phenology | Date of bud flush | BudFlush | ORL | 2009 |
|  |  |  | SAV | 2011 |
| Architecture | Branching angle | BrAngl | ORL | 2009 |
| Biochemical | H/G lignin ratio | H.G | ORL | 2011 |
|  |  |  | SAV | 2009 |
|  | S/G lignin ratio | S.G | ORL | 2011 |
|  |  |  | SAV | 2009 |
|  | Lignin content | Lignin | ORL | 2011 |
|  |  |  | SAV | 2009 |
|  | Glucose content | Glucose | ORL | 2011 |
|  |  |  | SAV | 2009 |
|  | Xylose to glucose ratio | Xyl.Glu | ORL | 2011 |
|  |  |  | SAV | 2009 |
|  | 5C/6C carbon sugar ratio | C5.C6 | ORL | 2011 |
|  |  |  | SAV | 2009 |
|  | Extractives content | Extractives | ORL | 2011 |
|  |  |  | SAV | 2009 |

**Figure 1:**
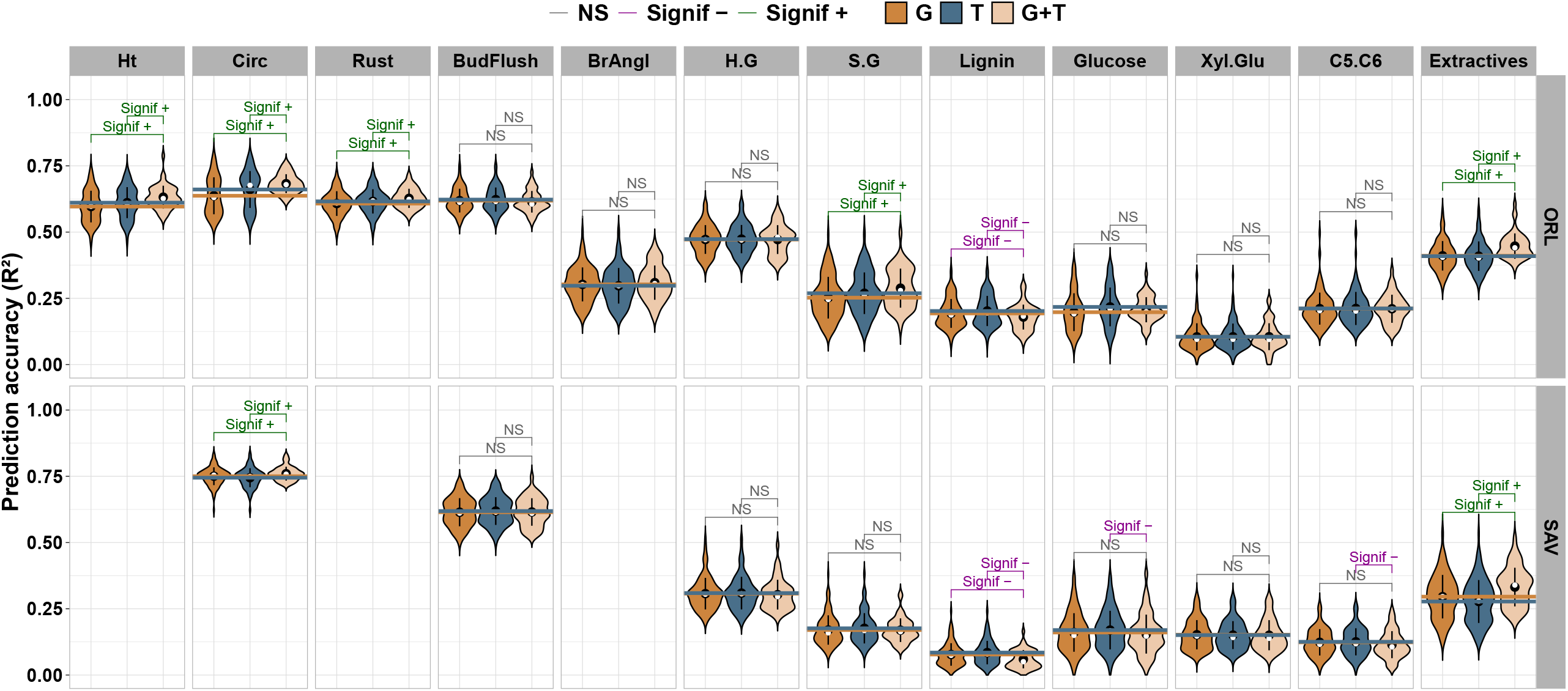
Prediction accuracies. Violin plots of prediction accuracies for 21 traits in the poplar dataset according to three models, using only genotyping data (the G model colored in dark brown to the left in the panels), using only transcriptomic data (T model colored in dark blue), and concatenating both genotyping and transcriptomic data (G+T model colored in light brown to the right). Each distribution of accuracies resulted from a cross-validation scheme. Significance from paired tests is shown for comparisons between models, with a sign indicating if the accuracy is increased (+) or decreased (-) in the multi-omics model by comparison with the single omic ones. Some traits were evaluated in two sites (“ORL” standing for Orléans in France and “SAV” for Savigliano in Italy). The white and black dots show the median and mean of the precision distributions, respectively. The dark brown and dark blue horizontal lines represent respectively the mean of precision distributions of G and T models.

We compared for each trait the prediction accuracies of both the single omic models with the multi-omics model, and tested if they significantly differed with a paired Wilcoxon signed-rank test. Over all traits, the differences between the average accuracy of multi-omics model compared to the single omic models ranged from -0.025 to 0.054. Seven of the 21 traits showed a significant gain with the multi-omics model over both the single omic models. These 7 traits included all the growth traits, the pathogen resistance trait, as well as 3 of the 14 biochemical traits (S.G_ORL, Extractives_ORL and Extractives_SAV). It is noteworthy that most of these traits (5/7) were measured in Orleans, the site where transcriptomics data were also collected. The only 2 traits presenting an advantage for the multi-omics model at the Italian site (Circ and Extractives) were also advantageous on the French site. Some traits showed a significant loss of accuracy with the multi-omics model, two (Lignin_ORL, Lignin_SAV) when the comparison was against the G counterpart, and four (Lignin_ORL, Lignin_SAV, Glucose_SAV and C5.C6_SAV) when it was with T model. Of note, all these traits displaying a decrease in accuracy with the multi-omics model were biochemical traits, they had low prediction accuracies and were more often than not measured in Savigliano (3 in Italy versus 1 in France).

In summary, the multi-omics model showed performance advantages over the single omic models in 7 of the 21 traits, more frequently on traits measured in Orleans where transcriptomic data were collected than in the Italian site. The multi-omics model underperformed the single omic models on 4 occasions, corresponding to 3 traits measured in Italy and one in Orleans. For the 10 remaining traits, no differences between models were detected (**Figure 1 and Supplemental Table S1**).

### eQTL analysis sheds light into the interplay between the genome and the transcriptome

To further gain insight into the interplay between the two omics layers for phenotype prediction, we studied their relationships through an eQTL analysis. Such analysis was performed with two specific detection steps, the first ignoring linkage disequilibrium between SNPs (called Step_0) and the second detecting multi-locus eQTLs (called Step_opt). The resulting eQTLs at both steps were further classified into CIS and TRANS regulatory elements according to their genomic proximity with the transcripts there were associated with (for more details see the method section). **Figure 2** and **Supplemental Figure S1** presents a map of these associations, respectively for step_opt and Step_0, with dot size reflecting the eQTL score. The darkened diagonal includes all CIS mediated associations, while the off-diagonal dots represent TRANS eQTLs. It is important to note that some positions at the marker axis present highly populated vertical trails across the genome, corresponding to important regulatory hubs.

**Figure 2:**
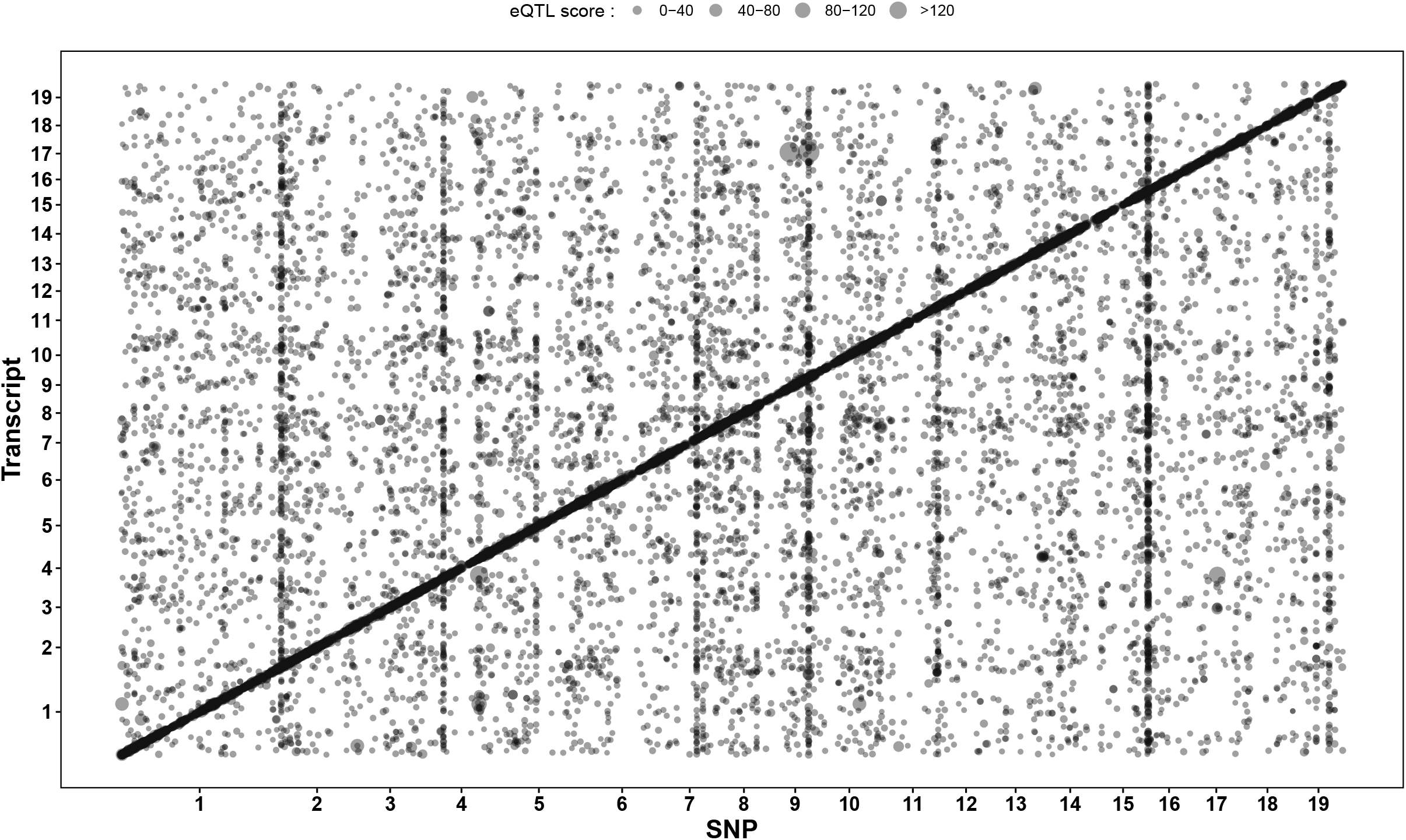
eQTL map between SNPs and transcripts. Map of associations (dots) between SNPs and transcripts through an eQTLs analysis with multi locus detection (Step_op), with dot size reflecting the association score (- log10 of the p-value of the test). The darkened diagonal includes all CIS mediated associations, while the off-diagonal dots represent the TRANS associations.

For both detection stages, we found eQTLs for 10,242 out of the 34,229 transcripts available in the transcriptomic dataset. Step_0 detected a total of 119,022 eQTLs on the marker dataset, including 72,841 (61.2%) CIS regulatory elements and 46,181 (38.8%) TRANS regulatory elements. At the optimal step of the eQTLs analysis (Step_opt), which accounted for linkage disequilibrium between SNPs, we detected a total of 18,248 eQTLs, of which 7,845 (43%) were CIS regulatory elements, and 10,403 (57%) were TRANS regulatory elements (**Supplemental Figure S2A**). CIS eQTLs displayed on average a larger effect than TRANS eQTLs (**Supplemental Figure S2B**). This point explains why there were so many at step_0, while their number drastically decreased at step_opt when LD was accounted for by the multi-locus approach. Whatever the step considered, the maximum distance between CIS eQTLs and their associated genes ranged between 12 and 14 kb.

### The importance of predictors is less preserved for traits displaying a predictive advantage with integration

To get insights into the factors explaining the gain or loss in predictive ability when using two omics by comparison with a single omic, we further looked at the variation in importance of the individual predictors over the two types of predictive models. For each of the three models, the importance of the different predictors was estimated as the rank of their squared effects from the ridge regression models.

We looked at correlations between the importance of predictors across single and multi-omics models, splitting the predictors into three categories determined from the eQTL analysis: TRANS eQTLs or regulated transcripts, CIS eQTLS or regulated transcripts and no eQTL. For SNPs, the correlation between the importances ranged from 0.62 to 0.99 across traits and predictor typologies. They were generally lower for the traits that also showed advantages with the G+T model over single omic models, and for those measured in the French site. Rust resistance, for instance, had the lowest correlations across the different categories among all measured traits (0.62, 0.63 and 0.65, respectively for TRANS eQTLs, CIS eQTLs and non eQTLS). Also, growth traits showed relatively low correlations compared to most of the traits, although this happened only for measurements in the French site (Ht_ORL, Circ_ORL), with those in the Italian site (Circ_SAV) being much higher and comparable to the top correlations. For the remaining traits, correlations between importances remained high, generally above 0.9 but with a few values close to 0.8 (**Supplemental Figure S3A**). The correlations between transcript importances (**Supplemental Figure S3B**) were generally lower than those for SNPs, varying between 0.52 and 0.89 across traits and predictor categories. Following a similar pattern as for SNPs, the traits showing the lowest correlations were also those for which the multi-omics displayed a predictive advantage over single omic models, as well as those measured in the French site. Growth and rust resistance traits were those showing the lowest correlations. Although with small differences, CIS-regulated transcripts showed lower correlations than those from TRANS-regulated counterparts, across traits and sites.

### TRANS-eQTLs show the most important changes of squared effect rank between multi-omics and single omic models

Previous correlations pinpointed to changes in importance in some of the categories of predictors. Such changes can be illustrated by the difference in importance (rank of squared effect) between the multi-omics model and that of the single omic model (either T or G, for transcripts and SNPs respectively) (see Methods for details).

When looking at the variation of the differences in importance (**Supplemental Figure S4, Figure S4**), the amounts were much larger for eQTLs (G+T versus G) than for targeted transcripts (G+T versus T). Higher variations were also found for TRANS eQTLs than for CIS counterparts, and for traits phenotyped at Orleans than for those in the Italian site. Thus, changes in importance occurred with more intensity for eQTLs, with a TRANS regulation, and linked to traits measured where the transcripts were sampled.

An alternative way of visualizing those changes is represented in **Figure 3**. This time, changes were averaged for a given trait and the resulting distribution of averages represented by predictor category and site. Patterns were very different between eQTLs and targeted transcripts, and also between sites. The most important changes in ranking happened at the Orleans site. With respect to predictor typologies, it was TRANS-eQTLs that showed the most important changes, with an overall loss of importance when switching to the G+T model, notably for the traits benefiting the most in performance from concatenation (growth and rust resistance). Less conspicuous were the changes for CIS-eQTLs, overall of negative sign but of lesser magnitude. Non-eQTLs showed generally small changes across traits. For targeted transcripts, the most impacted typology was CIS regulated genes, with an overall loss in ranking across traits, notably for growth and rust resistance traits.

**Figure 3:**
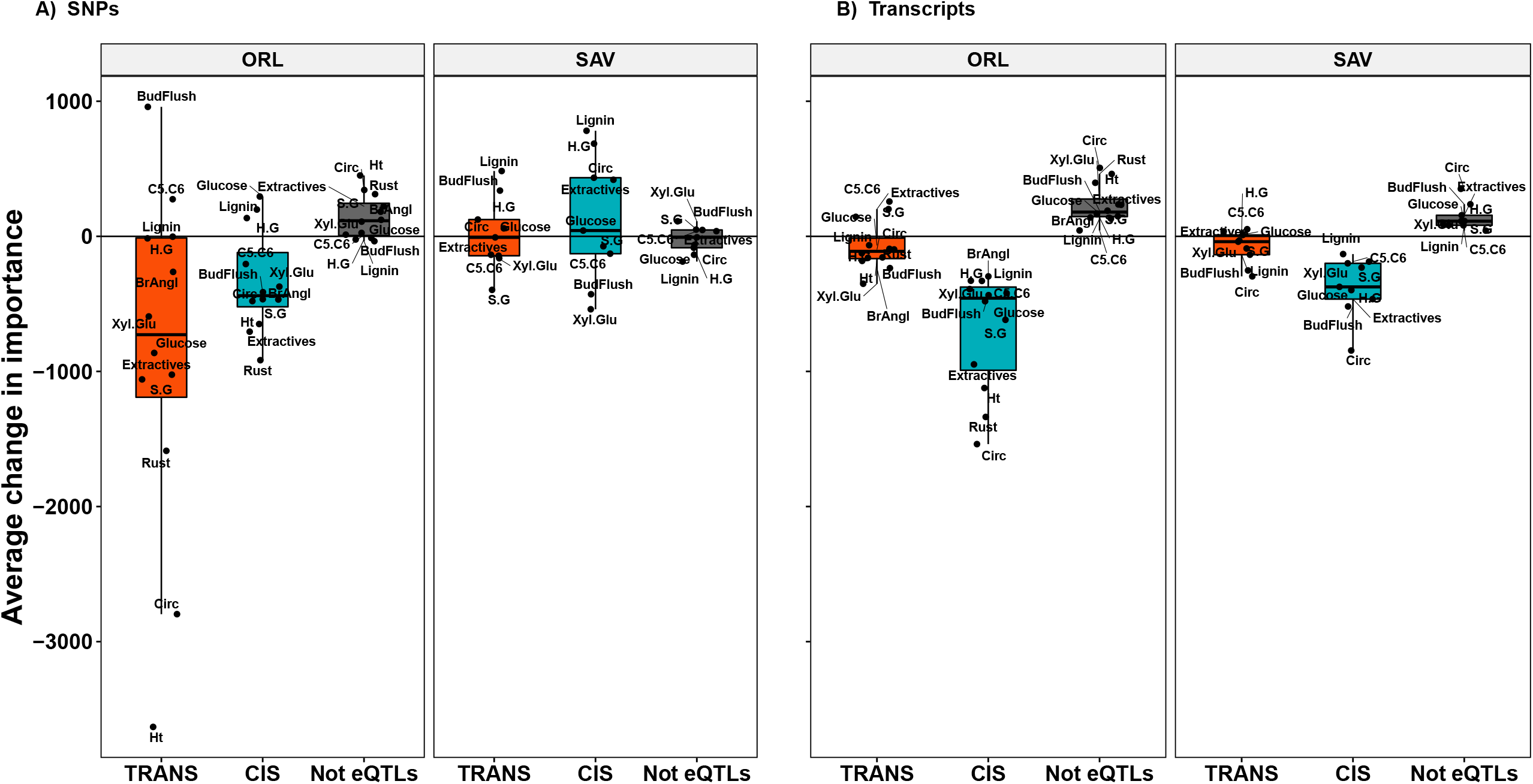
Distribution of predictors’ change in importance. Boxplot of the average change in importance of SNPs (panels A) and transcripts (panels B). Each dot represents the average difference per trait and per site of the predictor ranks between the multi omics model (G+T) and the single omic models (G for SNPs and T for transcripts). The red and blue boxplots show respectively the distribution of the average rank change for the TRANS-eQTLs and CIS-eQTLs. The boxplot in black shows the distribution for the predictors that have not been detected in the eQTL analysis.

### A negative relationship exits between the change in importance of eQTLs and CIS regulated transcripts and the predictive ability of the integrated multi-omic model

**Figure 4** represents the link across traits between average change in importance of predictors and advantage in performance of multi-omics over the single omic counterpart. In the case of eQTLs-TRANS, generally the most affected predictors following concatenation, a significant relationship (r=-0.81, p=0.0015) can be drawn where gains in prediction occurred at the expense of losses in ranking of predictors. A similar pattern, although of lesser magnitude, is to be found for eQTLs-CIS and CIS regulated genes (r=-0.6, p=0.037 and r=-0.64, p=0.024, respectively). No significant link was found for TRANS regulated genes. An equivalent representation for the Savigliano site showed no significant links across categories of predictors (**Supplemental Figure S6**).

**Figure 4:**
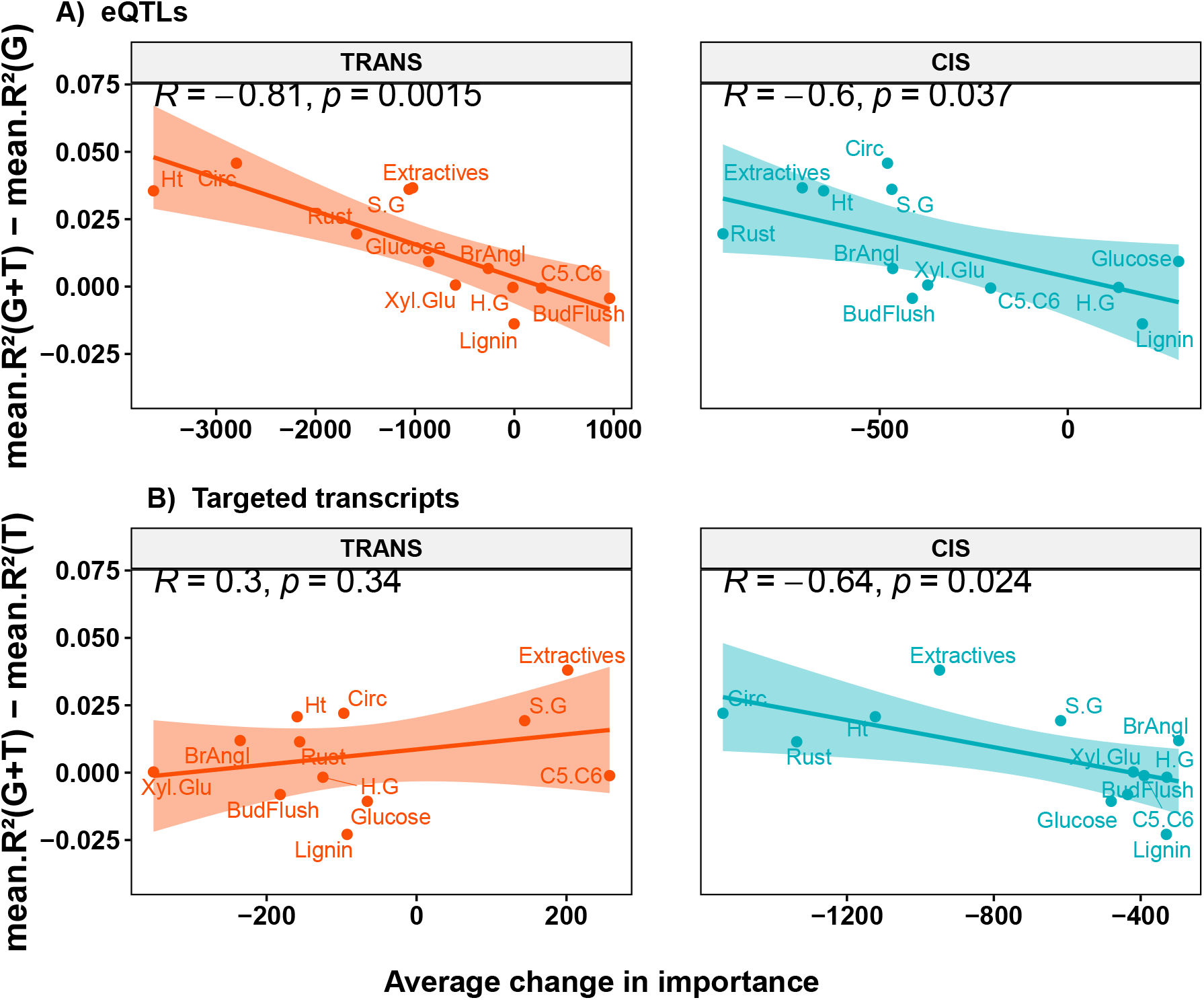
Relationship between predictors’ change in importance and muti-omics prediction advantage. Regression across traits measured at Orleans between average change in importance of predictors and advantage in performance of G+T over the single-omic counterpart. The top panel (A) shows the regression obtained with the eQTLs (TRANS-eQTLs on the left and CIS-eQTLs on the right). The bottom panel (B) shows the regression obtained with the regulated genes (TRANS on the left and CIS on the right).

### Gene ontology analysis suggests that top targeted transcripts or eQTLs are trait specific

We further selected the transcripts or eQTLs whose importance was most affected through data integration, by focusing on the 1% percentile at each extreme of the distributions, and carried an enrichment analysis in GO terms on the resulting features (**Figure 5 and Supplemental Table S2**).

**Figure 5:**
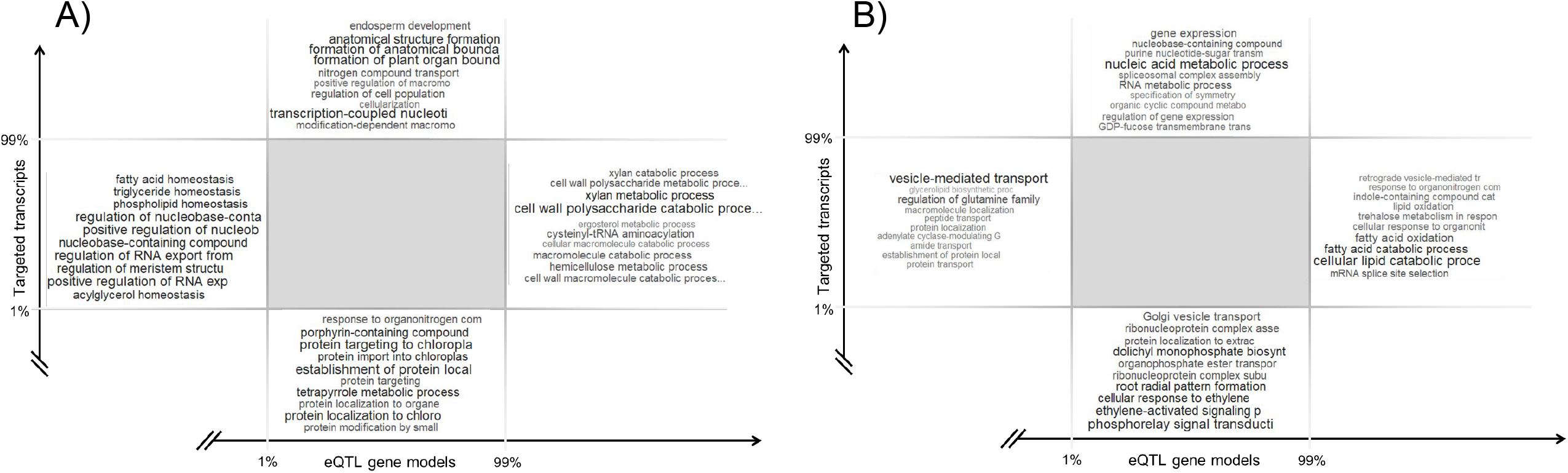
Gene ontology terms enrichment analysis. Schematic representation of the enriched GO terms among the top targeted transcripts or eQTL gene models list for A) the circumference of the tree trunk or B) the lignin content evaluated at Orleans. Font size and color intensity are proportional to -log10(p) of the top 10 GO terms.

For all type of traits, the analysis of the GOs showed enrichment of terms from general cell cycle process (e.g. “regulation of RNA export”, “regulation of nucleobase”, “positive regulation of RNA, vesicle-mediated transport”) in the lists of eQTL gene models selected as having the most negative impact on their importance during integration. The same results were visible for all traits with the lists of targeted transcripts selected with a negative effect, the GO terms enrichment were associated to ubiquity process like “protein targeting to chloroplast” or “protein localization to chloroplast” in the circumference of the tree trunk study (**Figure 5A**) or “phosphorelay signal transduction” and “cellular response to ethylene” for the lignin content study (**Figure 5B**).

For traits that showed significant gain with integration, this analysis suggests that the list of targeted genes with the most positive effects under integration are enriched with different terms specific to the traits. For example, we found cell wall related terms like “cell wall polysaccharide catabolic process”, “xylan metabolic process” for the circumference of the tree (**Figure 5A**). The same results were found for the eQTL gene models selected with the most positive effects with specific GO terms for tree circumference like “formation of plant organ” or “formation of anatomical boundary”.

On the contrary, in traits that showed significant predictive losses with multi-omics model over single omic ones, like the lignin content (**Figure 5B**), showed also enrichment in GO terms of general process for the most positive effect targeted transcripts or eQTL gene models (e.g. “nucleic acid metabolic process”, “gene expression” for the targeted transcripts and “cellular lipid catabolic”, “fatty acid catabolic process” for the eQTL gene models).

## Discussion

In this study, we used 21 traits to compare the relative advantages of integrating genomic and transcriptomic data for phenotype prediction versus using each omic separately. This relative advantage of integration over single omic varied across traits. For traits such as growth and pathogen resistance, integration yielded more accurate predictions than the single omic counterparts, while for most of the others, basically biochemical traits, no difference was detected, with still a few cases of underperforming concatenation. By using a simple modeling approach like ridge regression, we showed that gains in the traits benefited by integration were associated with systematic changes in importance for some specific predictors, and that those predictors were involved in the interplay between SNP polymorphisms and transcripts, pinpointing at adjustments in effects due to redundancies. Such findings at the statistical level were also backed up by a subsequent biological analysis of GO terms.

In order to better understand the reasons underlying trait differences in the benefits of concatenation, we sought to evaluate the interplay between the genomic and transcriptomic data, by making use of an eQTL analysis. Such analysis allowed us firstly to categorize the predictors into CIS eQTLs, TRANS eQTLs, non eQTLs, CIS regulated transcripts, TRANS regulated transcripts and transcripts with no eQTL detected. Secondly, based on such categorization, we could quantify the changes in predictor importance for each of these categories when using the multi-omics model by comparison with single omic ones. Over all the traits under study, we found a strong negative and significant correlation between the relative advantage of the multi-omics model compared to the single omic ones and the drop in importance of the predictors for eQTLs (R=-0.81 for TRANS eQTLs and R=-0.6 for CIS eQTLs) and transcripts regulated in CIS (R=-0.64). Such a relationship could be interpreted in terms of redundancy between predictors coming from different omics. Indeed, the traits that benefited the most from concatenation were also those for which TRANS eQTLs and CIS-regulated transcripts lost the most in terms of importance in the combined predictive model compared to the single omic counterparts, and thus for which the combined model decreased the redundancy between predictors by down weighting those that specifically matter for the eQTL versus transcript covariation. Redundancy, *per se*, would not necessarily explain gains or loss in performance, but down weighting redundant predictors could allow other minor predictors, otherwise silenced, climb in importance in such a way that the concatenation model improved in predictability.

To our knowledge, this study is the first to establish such a relationship between integration success and redundancy between omics layers, pinpointing eQTLs as key players in such interplay. It is worth mentioning that we could establish such a relationship because of the relatively large number of traits under study, compared to previous works (Ehsani et al., 2012; Guo et al., 2016; Morgante et al., 2020; Azodi et al., 2020; Li et al., 2019; Takagi et al., 2014; Schrag et al., 2018). The relative gains from integration ranged from -0.02 to +0.05 R^2^ across all 21 traits. These gains are indeed small, but are consistent with the state of the art. However, our objective here was to attempt to understand the factors that underlie this gain in order to produce new knowledge that will allow us to improve in more consequent ways the advantage of integration with other methods.

### A gain of integration was mainly found for traits related to growth and for traits evaluated in the same location as transcriptomic evaluation

We observed a significant advantage of multi-omics over single omic for all growth-related traits (Ht_ORL, Circ_ORL and Circ_SAV). Since growth results from cell division and expansion in the apical and cambial meristems (Xylem and cambium) (Chaffey et al., 2002) this relationship between the tissues from which we extracted transcripts and the growth traits (circumference and height) may explain the significant integration advantage observed for these traits.

The advantage of integration over the single omic models was also observed for leaf rust resistance. Although xylem and cambium, the tissues sampled for RNA sequencing, seem disconnected to a phenomenon occurring at the leaf level, the relationship here is likely indirect since links between resistance and growth have been reported by other studies (Wang and Kamp, 1992; Steenackers et al., 1996; Newcombe et al., 2001). Following the same reasoning, phenology and architectural traits considered here do not show clear relationships with cambial meristems or with growth-related traits, and therefore support the lack of benefits observed for them in the concatenation models.

For the majority of wood biochemical traits, the multi-omics integration model performed similarly or worse than the single omic model. It is noteworthy that these traits are overall not well predicted with single or multi-omics, and they originally come from near infrared spectroscopy prediction which may have included some noise to their variation. We could hypothesize that this factor underlies the observed poor performance for this category of traits during integration.

These series of observations across traits also point to the idea that transcripts capture some new information not necessarily available for SNPs, such as that associated to the genic interplays occurring at the specific tissue sampled for transcripts, which could be of a non-additive nature (gene-gene interactions), or even genotype-by-environment interaction effects, which are both not explicitly modelled when using exclusively SNPs as predictors.

Among the 21 analysed traits, we observed that the benefits of the concatenation model happened more often for the traits measured in Orleans where the transcriptomic data were also collected than for those in the Italian site. This advantage when phenotypic and transcriptomic data evaluation are carried out at the same location can be interpreted in terms of genotype-environment interactions effectively captured by the transcripts (Buil et al., 2015; Idaghdour and Awadalla, 2013). Conversely, for phenotypes evaluated at the Italian site, the transcriptomic data more likely bring redundant information to that of SNPs, which in turn do not result in any advantage in the multi-omic integration.

### Negative change in rank between the two models implies an increase in importance

Our goal was to identify factors that influence the success of genomic and transcriptomic data integration for phenotype prediction. To this end, we chose a simple integration method that allows us to track changes in the importance of each variable between the integration model and the single omic models. As described in Zampieri et al. (2019) and Ritchie et al. (2015), there are several ways to integrate multi-omics data, the simplest being the concatenation method. Using a ridge regression model, concatenation allows to directly estimate each predictor effect, accounting for all other variables (SNPs and transcripts), unlike LASSO and elastic-net where some degree of variable selection is applied, while trying to minimize the covariation between the predictors’ effects. This method allowed us to track the evolution of the relative importance of the predictors in the multi-omics model compared to the single models, and therefore infer potential redundancies by the changes in importance. Since the effects of SNPs or transcripts between the two models are not at the same scale to gauge importances, we had to bring them to a common scale. A simple and efficient way to do this is to work with ranks of the squared effects of predictors.

Comparing the changes in ranks between the multi-omics model and the single omic ones informed us about the gain or loss in importance of each predictor. A predictor will have a positive change in rank when it has high importance (low rank) under the single omic models and ends up with low importance (high rank) in the multi-omics model. Conversely, a negative change in rank between the two models implies an increase in importance. A zero rank change corresponds to a predictor that keeps the same importance between the two models.

### Integration success is driven by the loss in importance of covariation sources between genomics and transcriptomics

Our main hypothesis was that sources of redundancy between SNPs and transcripts play an important role in the success of the integration. The ideal candidates as a source of redundancy between SNPs and transcripts are eQTLs and the genes they regulate, so we performed an eQTL analysis to identify eQTLs (CIS and TRANS) and their regulated genes among our dataset. In order to remain in the same framework as in the ridge prediction models, we used the results of the eQTL analysis for which linkage disequilibrium was not taken into account, which enabled us to get information at the SNP level rather than at the locus level. However, the SNPs in our dataset are derived from RNAseq and are representative of the functional space of the genome, thus capturing few SNPs in the intergenic spaces. This might have affected our ability to detect some TRANS-eQTLs. Nevertheless, our multi-locus analysis showed that TRANS-eQTLs remained the majority, with some hotspot or hub loci associated with a fairly large number of transcripts. Such behavior has previously been reported in other species such as yeast (Albert et al., 2018) or maize (Liu et al., 2017; Swanson-Wagner et al., 2009).

The main results of our study is the strong negative and significant correlation found between the relative advantage of the multi-omic model over the single omic ones and the average importance losses of the eQTLs (more pronounced for TRANS) and the genes regulated in CIS. It is important to note here that CIS-regulated genes are on average regulated by more eQTLs than their TRANS-regulated counterparts, 10.15 and 6.63 eQTLs respectively (**Supplemental Figure S7**), suggesting that CIS-regulated genes are the source of more redundancies than TRANS-regulated genes in our dataset. To our knowledge, this study is the first to establish this direct relationship between integration success and losses in the importance of eQTLs and regulated genes. This relationship was possible to establish due to the relatively large number of diverse traits that we used.

There are results obtained in other studies that indirectly suggest the importance of eQTLs for multi-omics integration between genomics and transcriptomics. Ehsani et al. (2012) observed in mice losses in importance of eQTLs in the combined genomics and transcriptomics model versus the model with only genomics for the phenotype that shows an advantage of integration (body mass). Such behavior of eQTLs in this study was observed only for a single phenotype with low resolution genotyping data. Also, Ye et al. (2020) were successful in improving the performance of phenotype prediction in Drosophila using genotypes of eQTLs regulating genes that are important for the phenotype. They proceeded with successive selection steps involving a transcriptome wide association study (TWAS) with an eQTLs analysis for the TWAS significant genes, while optimizing the detection thresholds of these two analyses. Their results indirectly suggest the importance of eQTLs for the integration between genomics and transcriptomics. The negative correlation between the relative advantage of the multi-omics model over the single omic ones and the average losses in importance of covariation sources suggests that the integration success is driven by a minimization of the redundancy between genomics and transcriptomics. Azodi et al. (2020) observed in maize that concatenation between genomic and transcriptomic data improves the prediction of one of their 3 studied phenotypes. For this phenotype, they showed that the most important SNPs and transcripts were not redundant in the sense that they were not located in the same genomic regions, nor were they regulators of important transcripts.

### The observed redundancy may be explained by biological processes

Up to now, we have shown statistically that the sources of redundancy were penalized with a weaker importance under integration. Our GO enrichment analysis provided a more biological point of view to bring extra evidence of the role of redundancy. Generally speaking, genes pointing at general ubiquitous biological processes were more likely the source of redundancy, while those associated with specific processes could more easily bring extra useful information to the prediction process. The GO analysis showed that genes gaining in prediction importance under integration were generally associated with specialized processes of relevance for the predicted phenotype. This pattern was observed notably for traits related to wood production. On the contrary, the genes that were most heavily affected in their importance under integration showed a characteristic enrichment of terms linked to the general cell cycle processes. As the transcriptomic data came from young differentiating xylem and cambium tissues, the redundancy (and complementarity) that we observed is strongly associated with phenotypes related to the production of wood, like the trunk circumference. This interpretation of our results might also apply to the loss of prediction for traits whose genes are not likely to be represented in our transcriptome, such as those related to rust resistance for example. One eventual validation could be to complement the transcriptomic data with alternative collection on tissues other than those closely connected to xylem and cambium, like leaves, and work on traits more specifically expressed on the collected tissue (i.e. rust resistance). If the genes associated with the general biological processes are found to be sources of redundancy through GO analysis, one strategy to improve prediction could be to reduce or minimize their contribution to the models.

### Perspectives

One of the main findings of this study is the fact that certain predictors with ubiquitous connections seem to be made redundant when integration takes place, leaving the stage for other features to be picked up, eventually less prominent but bringing true complementarity to the integrative prediction. For the sake of simplicity, our study could not take the extra step to devise a novel alternative to account for such redundancies. However, it would be quite straightforward to outline a basic strategy where the importance of predictors going into the model is penalized according to some function describing their redundancy in the data. Under kernel-based integration, for instance, some kind of optimization of composition in features included in the relatedness matrices could be devised so that the resulting kernels bring complementary information. Under a model-based integration, a multistage approach could be devised where associations between all involved omics are firstly carried out, so that the features contributing the most to the associations can be subsequently penalized to some degree or filtered out when it comes to construct a consensual model. More research would be required to devise and test a strategy to derive robust weightings.

It would be essential to gather extra information on the beneficial role of multiple omics, collected at different development stages or distinct tissues, so that links to different traits can be drawn. This is certainly a costly endeavor, which could be focused on specific training populations. Ideally, integration studies on those training sets would allow us to identify important hubs in the genetic architecture of traits, and use that information for differential weighting on other related populations with no or basic access to extra omics layers.

## Material and methods

### Plant material, experimental design and phenotypic evaluation

We studied 241 genotypes of *Populus nigra* originated from 11 major river catchments across 4 countries and representative of the species range in Western Europe. These poplars were evaluated in common garden experiments located on 2 contrasting sites (Orleans noted ORL and Savigliano noted SAV) (Guet et al., 2015). In each site, the experimental design consisted in a randomized complete block design with 6 blocks, and thus 6 repetitions per genotype. Twelve traits were evaluated on the 2 sites, as previously described (Gebreselassie et al., 2017; Chateigner et al., 2020). We considered traits measured in 2 sites as different traits, leading to a total of 21 traits (detailed in **Table 1**). These traits can be categorized into 5 types: growth, pathogen tolerance, phenology, architecture, and biochemistry. At Orleans, the trees were grown through 3 successive cycles: 2008-2009, 2010-2011 and 2012-2015. During the first growth cycle (2009), rust tolerance (Rust) was measured with a discrete score from 1 (no symptom) to 8 (generalized symptoms), as detailed in Legionnet et al. (1999). Average branch angle (BrAngl) was evaluated with a score on proleptic shoots from 1 to 4 (score 1: between 0° and 30°; score 2: between 30° and 40°; score 3: between 40° and 55°; score 4: and between 55°and 90°). During the second growth cycle, height (Ht) and circumference at 1-meter above the ground (Circ) were measured on 2 year-old trees (winter 2011). At Savigliano, trees went through two cycles: 2008-2009 and 2009-2010. Only Circ was measured during the second growth cycle on 2 year-old trees (winter 2010). Biochemical traits consisted in predictions of several chemical compounds obtained from near-infrared spectra on wood samples collected in the same years as growth traits and at both sites, as described in Gebreselassie et al. (2017). Biochemical traits included: extractives content (Extractives), total lignin content (Lignin), ratios between different lignin components like p-hydroxyphenyl (H), guaiacyl (G) and syringyl (S) (H.G, S.G), total Glucose content (Glucose), ratio between Xylose and Glucose content (XylGlu) and the ratio between 5 and 6 carbon sugars (C5.C6). One phenological trait was also measured, BudFlush as discrete scores for a given day of the year, measured on the apical bud (Dillen et al., 2009).

### Phenotype adjustments

All 21 traits were independently adjusted to field micro-environmental heterogeneity using the breedR package (Munoz and Sanchez, 2017). The model included blocks and spatial effects (autoregressive residuals function) to account for micro-environmental heterogeneity. Also a model selection was carried out using the AIC to select the effects to be included in the model and to tune the autoregressive parameters. The genotypic adjusted means from these models were used as phenotype for this study.

### Genotype and transcriptomic data

RNA sequencing was carried out in 2015 on young differentiating xylem and cambium tissues collected on two replicates of the 241 genotypes located into two blocks of the Orleans design (Chateigner et al., 2020). We obtained sequencing reads for 459 samples corresponding to 218 genotypes with two replicates and 23 genotypes with 1 replicate. These sequencing reads were used to provide both transcriptomic and genomic data.

For transcriptomic data, the reads were mapped on the *Populus trichocarpa* v3.0 primary transcripts and read counts were retrieved for 41,335 transcripts. Only transcripts with at least 1 count in 10% of the individuals were kept, yielding 34,229 features. The raw count data were normalized by Trimmed Mean of M-values using the R package edgeR v3.26.4 (Robinson and Oshlack, 2010) and we calculated the counts per millions (Law et al., 2014). To make the CPM data fit a Gaussian distribution, we computed a *log*2(*n*+1) instead of a *log*2(n+0.5) typically used in a voom analysis (Law et al., 2014), to avoid negative values, which are problematic for the rest of the analysis. For each transcript the log2(n+1) of the CPM were fitted with a mixed model including experimental (batch) and genetic effects to extract their genotypic blups. Those transcripts’ genotypic blups were used for the rest of our analysis.

The genotyping data was obtained, first by mapping the RNAseq reads on the *P. trichocarpa* genome reference (v3.0) (Goodstein et al., 2012). After the mapping, the SNPs were called using 4 callers. In order to generate a high-confidence SNP set we selected only the SNPs identified by at least 3 of the 4 callers and with less than 50% of missing values. Remaining missing values were imputed using complementary genotyping data obtained with a 12k Illumina Infinium Bead-Chip array (Faivre-Rampant et al., 2016). Full details of SNP discovery, data filtering criteria and final selection are given in Rogier et al. (2018). We then detected 874,923 SNPs. From these detected SNPs 428,836 SNPs were retained for this study after filtering the SNPs with minimum alleles frequencies lower than 0.05.

### eQTLs Analysis

eQTLs analysis was performed using the Multi-Loci Mixed-Model (MLMM) approach (Segura et al., 2012) and implemented in the R package MLMM v0.1.1. MLMM uses a step-by-step forward inclusion and backward elimination approach under a mixed-model framework which accounts for the confounding usually attributed to population structure with a random polygenic effect. For each of the 34,229 transcripts we ran MLMM for up to 10 steps and identified the optimal model according to the mBonf criterion (all selected SNPs are significant at a 5% Bonferroni corrected threshold). The initial and the optimal steps outputs have been saved for further analyses.

Based on the positional proximity of the genes, the eQTLS detected at each of these 2 steps were classified as CIS regulatory elements (non-coding DNA regulating the transcription of neighboring genes), and/or as TRANS regulatory elements (regulating the transcription of distant genes), according to the following rules:

- all eQTLs associated with the expression of a gene located in a different chromosome are classified as TRANS, and the targeted gene is classified as a TRANS regulated gene;
- all eQTLs located on the same locus as the gene it targets, according to the genome annotation, are classified as CIS, and the targeted gene is also classified as CIS regulated gene;
- the remaining eQTLs whose target gene is on the same chromosome but not on the same locus, were splitted into CIS or TRANS according to their distance to the middle of the gene they target. We estimated the maximum distance between the CIS eQTLs identified at previous step and the middle of the gene they target as 18.9 kb (eQTL being on the same position as its target gene). If the distance between eQTLs and the gene they target is greater than 18.9 kb they were classified as eQTLs TRANS and target Gene TRANS. Otherwise, the eQTLs and the target gene were classified as CIS.

### Models, prediction accuracy and cross-validation

Two ridge regression models were built for each trait with a single omic data as predictor, genotypic data (**G** model), or transcriptomic data (**T** model), respectively with p = 428,836 SNPs and q = 34,229 transcripts’ expression levels variables. A third multiomic was also built with integration by concatenation of both omics data (**G+T** model). These 3 models can be written as:

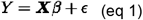

Respectively for models **G**,**T** and **G+T, X** represent the genotyping matrix(***n*×*p***), the transcript expression level matrix for the genes(***n*×*q***) and the concatenated transcript expression level and genotyping matrix(***n*× (*p*+ *q***)). With the same logic, ***β*** represent the vector of effect sizes of variables of those matrices. Y is the vector of phenotype, and the vector of residual errors of the model.

The models were computed using the R package glmnet (Friedman et al., 2010) in a 5 inner-fold and 10 outer-fold nested cross validation framework (Varma and Simon, 2006). The sampling process for the different folds was repeated 50 times. Each cross-validation sample was used across all traits and for the three models. Paired t-tests in R (rstatix package version 0.7.0) (Kassambara, 2021) were used for model comparisons of performance.

The models performances was measured using ***R***^**2**^ between observed and predicted values.

### SNPs and transcript effects ranking

In order to study the changes operated for each feature (SNP or gene) when changing prediction models from single omics to the concatenated counterpart, we compared the change in ranking of the effects across models. Ranks were obtained for each predicting model and trait from the ordering of squared effect sizes.

For each variant typologies, the estimated effects rank was compared between the single omic models (**G** or **T**) and the multi-omics model with a paired wilcoxon test and a Pearson correlation.

The difference in effect ranking between the model with concatenation and the single omic models was calculated for the different typologies of each feature (SNPs and genes). Then this ranking difference was averaged for each trait and regressed with the concatenation advantage of each trait, which is the average accuracy difference between the concatenation model and the single omics models:

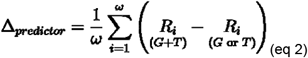

Predictor represents either SNPs or transcripts. R is the ranking vector of squared effect sizes of the given predictor. ***ω*** is the number of predictors (p for SNPs and q for transcripts). **Δ**_***predictor***_ is the average difference in effect ranking between the multi-omics model and the single omic one by trait for the given predictor.

### GO analysis

Functional enrichment was conducted based on the gene ontology (GO) terms associated with the best *Arabidopsis thaliana* homolog and based on Phytozome v12.1.6 database (Goodstein et al., 2012). GO analysis was conducted using R package topGO 2.44.0 (Alexa and Rahnenfuhrer, 2021) and Fisher’s exact test with ‘elim’ used to correct for multiple comparisons. The significant threshold of GO terms was *P*□≤□0.05.

## Availability of data and code

Supporting data is available at: https://doi.org/10.15454/8DQXK5

Code for running the test and replicate the analysis are available at: https://github.com/Tawfekh/Code-Article-Multi-omics-prediction

## Acknowledgements

The authors gratefully acknowledge the staff of the INRAE GBFOR experimental unit for the establishment and management of the poplar experimental design in Orléans, the collection of wood samples in each site, and their contribution to phenotypic measurements on poplars in Orléans; Alasia Franco Vivai staff for management of the poplar experimental plantation in Savigliano, and M. Sabatti and F. Fabbrini for their contribution to phenotypic measurements on poplars in Savigliano. We acknowledge the staff of BioForA for their contribution to RNA collection in the field. We also acknowledge the platform POPS for their implication with RNA sequencing and sequence analyses and Odile Roger for her contribution to sequence analyses.

Establishment and management of the experimental sites were carried out with financial support from the NOVELTREE project (EU-FP7-211868). RNA collection, extraction, and sequencing were supported by the SYBIOPOP project (ANR-13-JSV6-0001) funded by the French National Research Agency (ANR). This study was funded by the European Union Horizon 2020 research and innovation programme under grant agreement No 773383 (B4EST project) and by INRAE SELGEN metaprogramme (EPINET project).

## Author contributions

ARW: analyzed all the data and wrote the manuscript; HD: performed GO analyzes and wrote the manuscript; LS and VS: planned and designed the project, supervised the research and wrote the manuscript.

## Supplemental Data

**Supplemental Figure S1:** eQTL map between SNPs and transcripts (Step_0)

**Supplemental Figure S2:** Abundance and score of CIS and TRANS eQTLs

**Supplemental Figure S3:** Comparison between the importance of predictors across single and multi-omics models

**Supplemental Figure S4:** Variation of the change in importance of the eQTLs and targeted transcripts

**Supplemental Figure S5:** Change in importance of the eQTLs and their corresponding targeted transcripts

**Supplemental Figure S6:** Relationship between predictors change in importance and muti-omics prediction advantage for traits measured at Savigliano

**Supplemental Figure S7:** Average number of connection for the eQTLs and targeted transcripts

**Supplemental Table S1:** Prediction accuracies comparison between the multi-omics model and the single omic ones

**Supplemental Table S2:** Complete gene ontology analysis of all traits

## Notes

https://doi.org/10.15454/8DQXK5

https://github.com/Tawfekh/Code-Article-Multi-omics-prediction

### Competing Interest Statement

The authors have declared no competing interest.

