## Supplementary material for "eQTLs are key players in the integration of genomic and transcriptomic data for phenotype prediction": Supp Figure 1

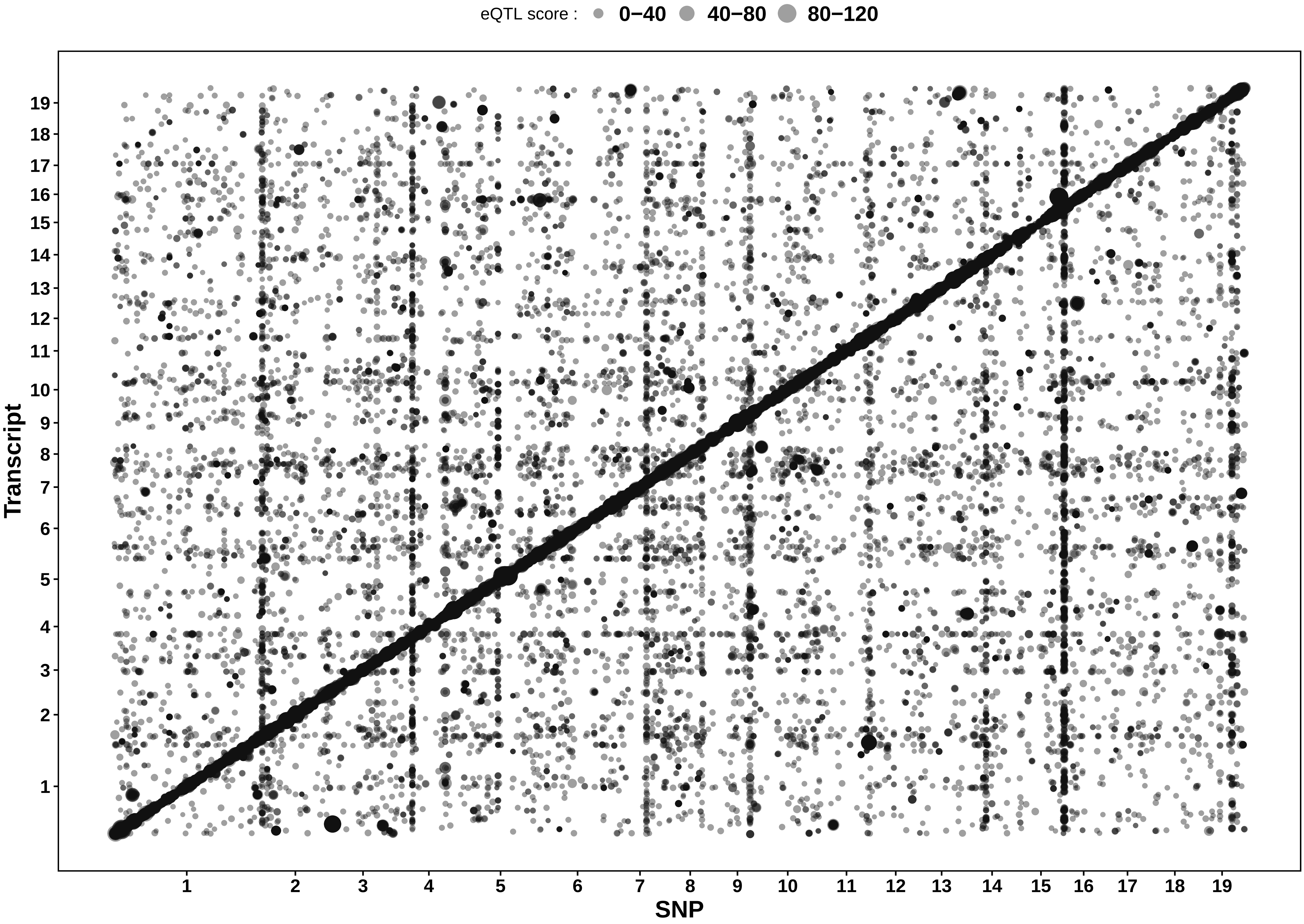


**Supplemental Figure S1: eQTL Map between SNPs and transcripts (Step_0)**

Map of associations (dots) between eQTLs and transcripts through an eQTLs analysis without taking into account the linkage disequilibrium between SNPs (Step_0).The dot size reflects the eQTL score. The darkened diagonal includes all CIS mediated associations, while the off−diagonal dots represent the TRANS associations.
