## Supplementary material for "eQTLs are key players in the integration of genomic and transcriptomic data for phenotype prediction": Supp Figure 2

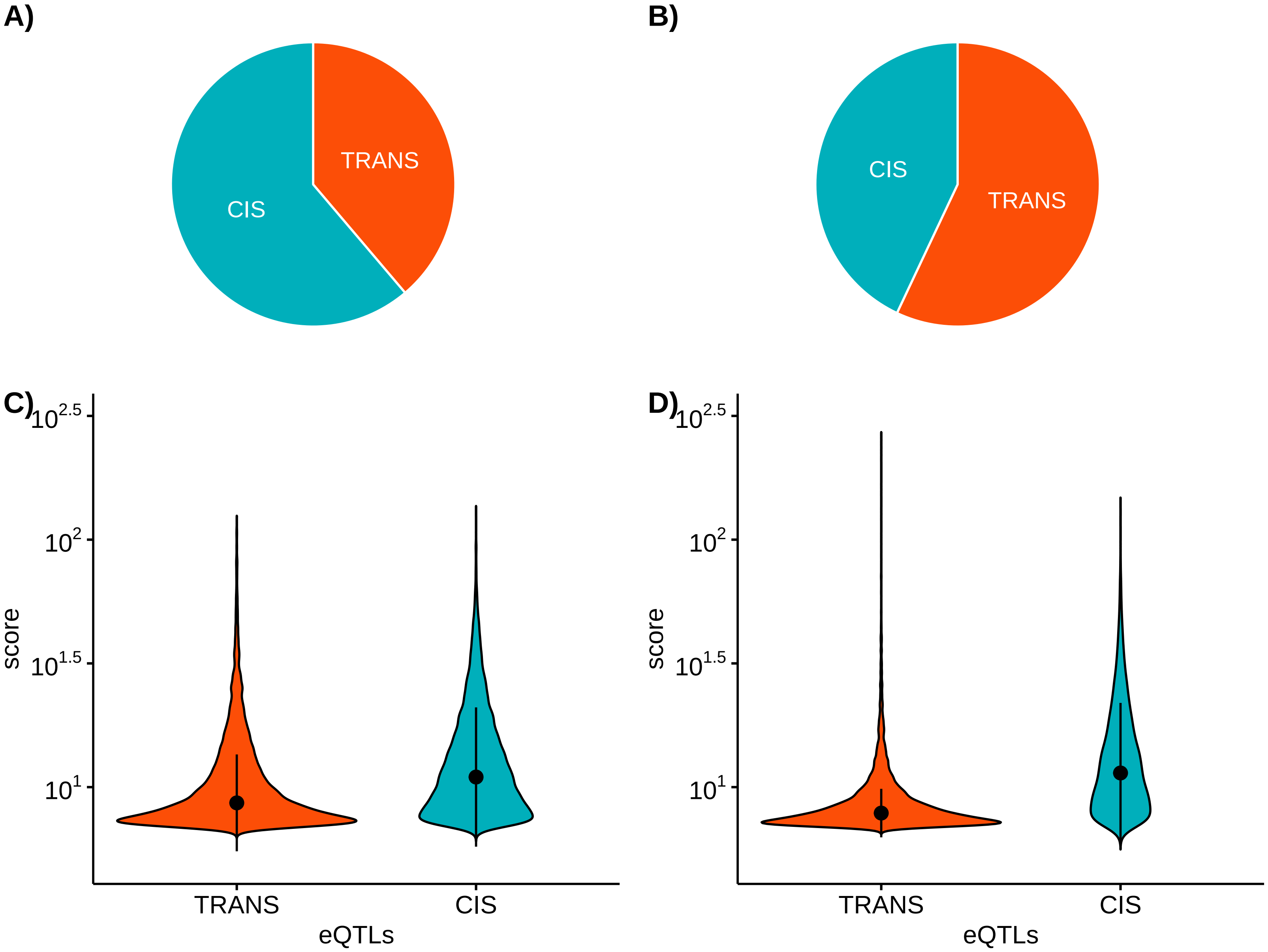


**Supplemental Figure S2: Abundance and score of CIS and TRANS eQTLs**

Plots A and B show respectively the proportions of CIS and TRANS eQTLs for the step of the eQTLs analysis without taking into account the linkage disequilibrium (step_0) and the optimal step that takes into account the multi−locus detection (step_op ). Plots C and D show respectively the detection scores of the CIS and TRANS eQTLs for the step_0 and step_op.
