## Supplementary material for "eQTLs are key players in the integration of genomic and transcriptomic data for phenotype prediction": Supp Figure 3

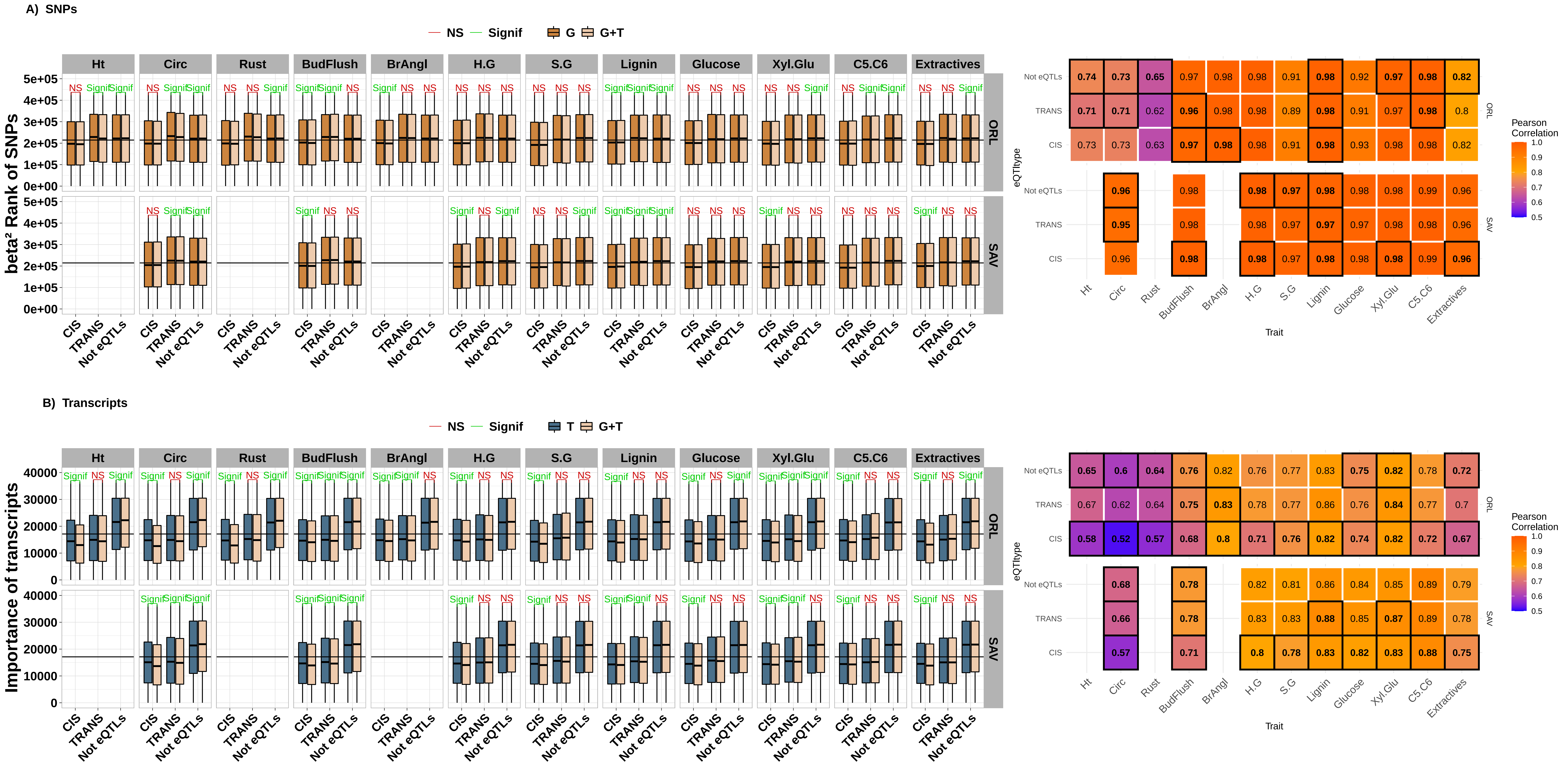


### Supplemental Figure S3: Comparison between the importance of predictors across single and multi−omics models

Top left and bottom left represent respectively the boxplots of SNPs and Transcripts importances, according to the traits and sites. Significance from paired tests is shown for comparisons each predictor importance between the multi omics models and the single ones, according to  the type of regulations. Top right (SNPs) and bottom right (transpcripts) represente the correlations between the importance of predictors across single and multi−omics models, splitting the predictors into three categories determined from the eQTL analysis: TRANS eQTLs or regulated transcripts, CIS eQTLS or regulated transcripts and no eQTL.
