## Supplementary material for "eQTLs are key players in the integration of genomic and transcriptomic data for phenotype prediction": Supp Figure 4

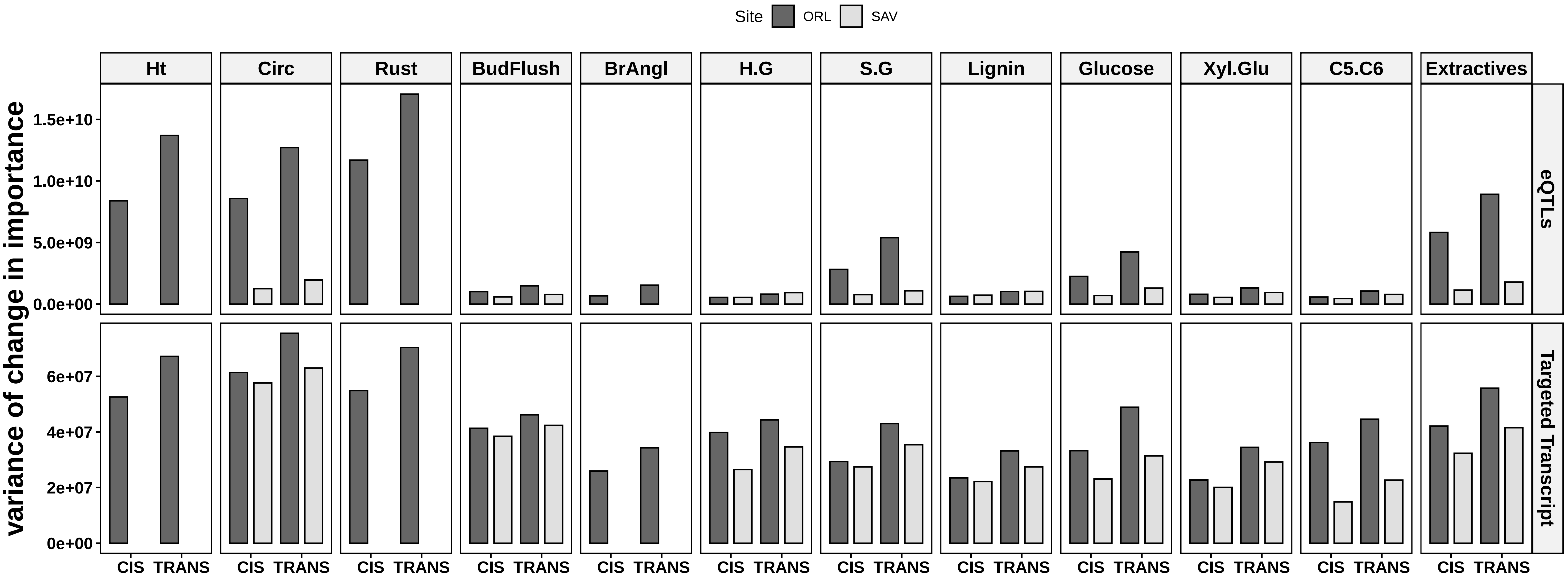


**Supplemental Figure S4: Variation of the change in importance of the eQTLs and targeted transcripts**

Barplot of the change in importance variance of eQTLs (top) and targeted transcripts (bottom) for each trait(each panels) and site(dark grey: Orléans ; light grey: Savigliano).
