## Supplementary material for "eQTLs are key players in the integration of genomic and transcriptomic data for phenotype prediction": Supp Figure 5

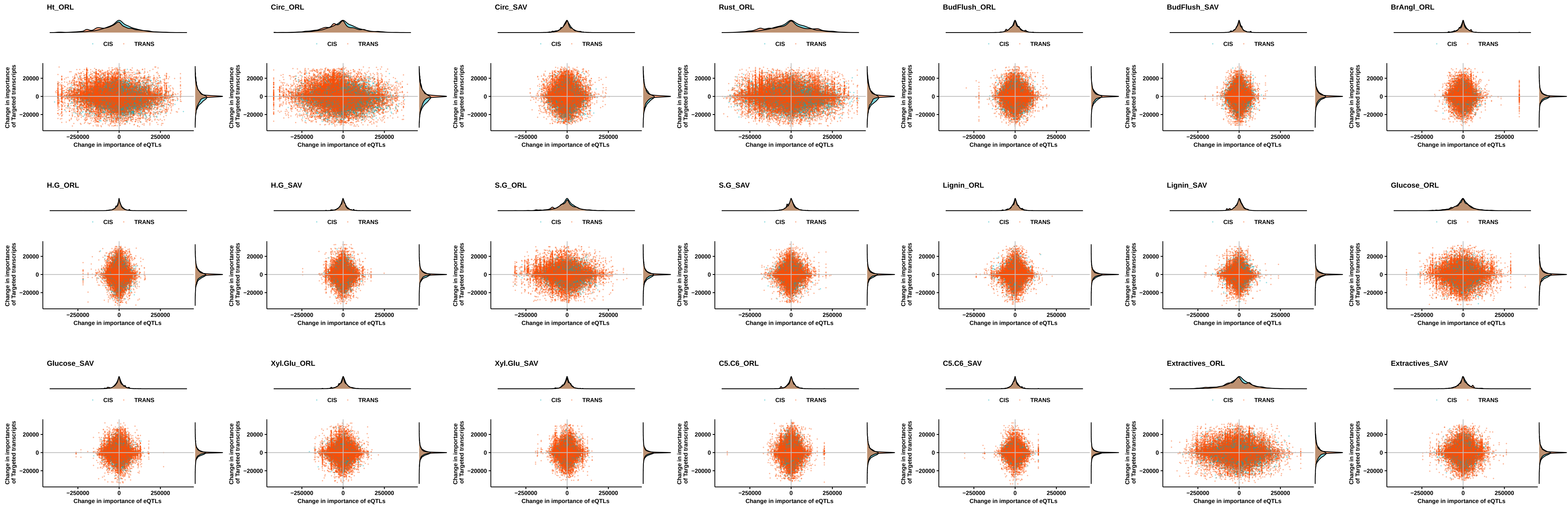


**Supplemental Figure S5: Change in importance of the eQTLs and their corresponding targeted transcripts**

Scatter plot of the changes in importance of the eQTLs and their corresponding targeted transcripts for each trait. The red and blue spots represent the regulation in TRANS and CIS, respectively.
