## Supplementary material for "eQTLs are key players in the integration of genomic and transcriptomic data for phenotype prediction": Supp Figure 6

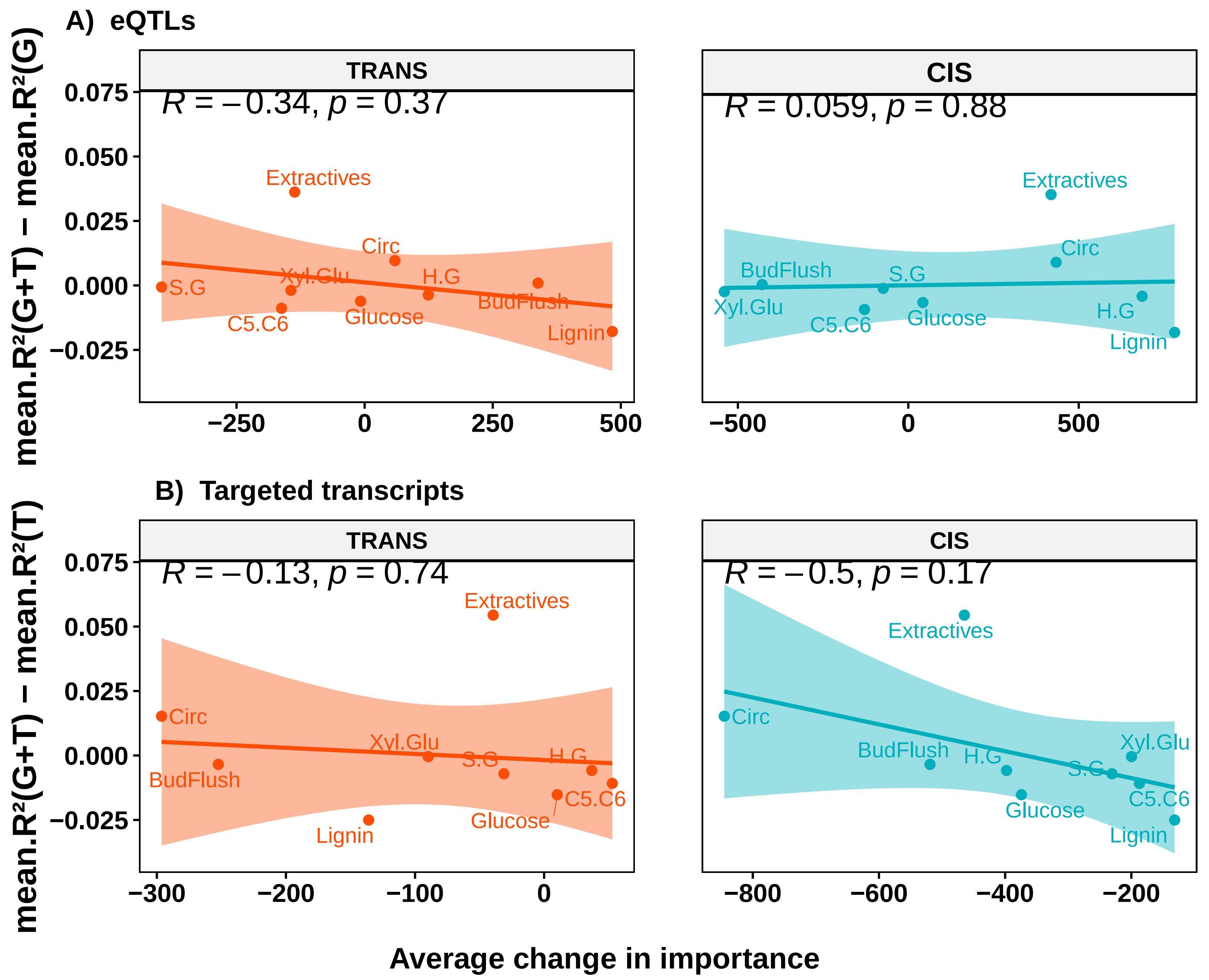


**Supplemental Figure S6: Relation between predictors change in importance and muti−omics prediction advantage for traits measured at Savigliano**

Regression across traits measured at Savigliano between average change in importance of predictors and advantage in performance of G+T over the single−omic counterpart. The top panel (A) shows the regression obtained with the eQTLs (eQTLs Trans on the left and eQTLs CIS on the right). The bottom panel (B) shows the regression obtained with the regulated genes (Trans on the left and CIS on the right).
