## Supplementary material for "eQTLs are key players in the integration of genomic and transcriptomic data for phenotype prediction": Supp Figure 7

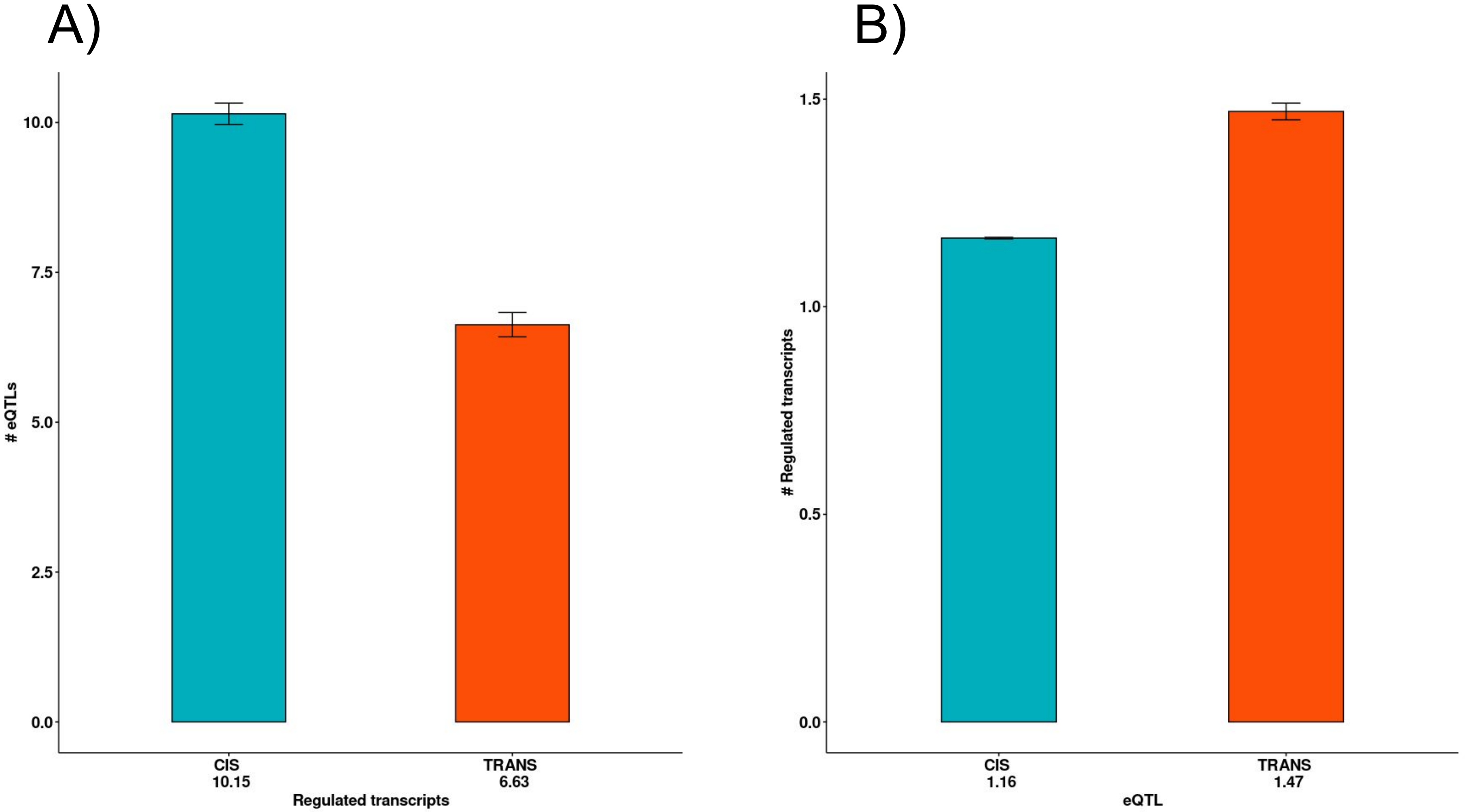


**Supplemental Figure S7: Average number of associations for the targeted transcripts and eQTLs**

Plots A and B show respectively the average number of associations (step_0) for targeted transcripts and eQTLs. The error bars indicate the standard errors.
